# Significant and conflicting correlation of IL-9 with *Prevotella* and *Bacteroides* in human colorectal cancer

**DOI:** 10.1101/2020.04.28.066001

**Authors:** E Niccolai, E Russo, S Baldi, F Ricci, G Nannini, M Pedone, FC Stingo, A Taddei, MN Ringressi, P Bechi, A Mengoni, R Fani, G Bacci, C Fagorzi, C Chiellini, D Prisco, M Ramazzotti, A Amedei

## Abstract

**Background:** Colorectal cancer (CRC) is a widespread disease that represents an example of chronic inflammation-associated tumor. In fact, the immune system, besides protecting the host from developing tumors, can support the CRC progression. In this scenario, the gut microbiota (GM) is essential to modulate immune responses and a dysbiotic condition can favor chronic/abnormal immune activation that support the tumor growth. GM can elicit the production of cytokines, influencing the immunostimulatory or immunosuppressive reactions, such as the tendency to mount Th1, Th17, Tregs or Th9 responses that play different roles towards colon cancer. Paradigmatic is the role of IL-9 that can both promote tumor progression in hematological malignancies and inhibit tumorigenesis in solid cancers. Therefore, to investigate the microbiota-immunity axis in CRC patients is crucial to well understand the cancer development with positive relapses in prevention and treatment.

**Aim:** The cellular and molecular characterization of the immune response and the evaluation of GM composition in healthy and tumor mucosa, focusing on the correlation between cytokines’ profile and GM signature.

**Methods:** We collected tumoral (CRC) and healthy (CRC-S) mucosa samples of 45 CRC patients. For each sample, we characterized the Tissue Infiltrating Lymphocytes (TIL)’s subset profile and the GM composition. In addition, in 14 CRC patients, we evaluated the CRC and CRC-S molecular inflammatory response (26 cytokines/chemokines) and we correlated this profile with GM composition using the Dirichlet Multinomial Regression.

**Results:** The analysis of T cells subsets distribution showed that CRC samples displayed higher percentages of Th17, Th2, Tregs, Tc17, Tc1/Tc17, and Tcreg, compared to CRC-S. Notably, also the number of Th9 was higher, even if not significantly, in CRC tissue compared to healthy one. In addition, we found that MIP-1α, IL-1β, IL-2, IP-10, IL-6, IL-8, IL-17A, IFN-γ, TNF-α, MCP-1, IL-1α, P-selectin and IL-9 were significantly increased in CRC compared to CRC-S. Moreover, the GM analysis revealed that CRC samples had significantly higher levels of *Fusobacteria*, *Proteobacteria*, *Fusobacterium*, *Ruminococcus2* (*Lachnospiraceae* family) and *Ruminococcus* (*Ruminococcaceae* family) than CRC-S. Finally, we found that the abundance of *Prevotella spp* in CRC samples was negatively correlated with IL-17A and positively with IL-9. In addition, the abundance of *Bacteroides* and *Escherichia/Shigella* species in CRC samples showed a negative association with IL-9 and IP-10 respectively.

**Conclusions:** Our data show a clear dissimilarity of inflammatory profile and GM composition between the tumor and the adjacent healthy tissue, displaying the generation of a peculiar CRC microenvironment. Interestingly, relating the tissue cytokine profile with the GM composition, we confirmed the presence of a bidirectional crosstalk between the immune response and the host’s commensal microorganisms; in detail, we documented for the first time that *Prevotella spp.* and *Bacteroides spp.* are correlated (positively and negatively, respectively) with the IL-9, whose role in CRC development is still debated.

## 1. INTRODUCTION

Colorectal cancer (CRC) is a complex and widespread disease, being the second cause of cancer-related deaths in the world^1^. Usually it begins as benign polyps that can become (especially the adenomatous type) cancerous if not removed. In human, only 10% of adenomatous polyps evolve in cancer; since the multi-step colorectal tumorigenesis does not involve exclusively genetic factors but is also deeply influenced by host factors, such as inflammatory and immune responses^2 3^. Indeed, chronic inflammation increases cancer risk through a deregulated activation of the immune system, which cause the loss of tissue architecture and genotoxic cellular DNA damages^4^. CRC is one of the best examples of chronic inflammation-associated tumor, occurring often in patients with inflammatory bowel disease^5^. Moreover, according to the immunoediting theory, the adaptive immune system, besides protecting the host from developing tumors^6^, can support the tumor progression. Specifically, T cells can develop different functional features during the cancer growth, affecting the disease progression and/or regression. The protective immunity is mediated by effector cells (Th1, and Th17/Th1) while “not effector” T lymphocytes’ subsets (Th2, Tregs, Tnull) can promote colon cancer progression^7 8 9^. In this scenario, also the microbiota plays an important role, since it is essential to modulate immune responses favoring the equilibrium between protective immunity and tolerance to commensal bacteria^10^. A perturbation of the gut microbiota (GM) composition can disrupt this balanced ecosystem, determining a chronic/abnormal immune activation and supporting the tumor growth. In fact, over the past 10 years, both specific bacteria and dysbiotic conditions have been associated or implicated in colorectal carcinogenesis^11 12^, sometimes through the interaction with the immune system ^13^. In particular, the role of *Fusobacterium nucleatum* is paradigmatic because it promotes the CRC by either the induction of epithelial cell proliferation^14^, thus generating a proinflammatory microenvironment propitious to cancer progression^15^ or the production of proteins able to block the cytotoxic anti-tumoral activity of T and NK cells^16 17^.

Moreover, the microbes can affect cancer cell antigenicity and adjuvanticity^18^, determining whether an antigen triggers an immune response and if its nature drive the acquisition of a specific T cell phenotype (effector or regulator). In addition, microbiota can elicit the production of cytokines (and other immune mediators), influencing the immunostimulatory or immunosuppressive reactions, such as the tendency to mount Th1/Tc1 (characterized by IFN-γ production), Th2/Tc2 (IL-4 and IL-13), Th17/Tc17 (IL-17) or Th9 (IL-9) responses^19 20 21^, that play different roles towards colon cancer ^9 22^. For example, the commensal bacteria can stimulate the *lamina propria’s* dendritic cells to produce IL-6, TGF-β and IL-23 needed to elicit the Th17 and Th9 lymphocytes development23, which have a dual role in CRC promotion^7 24^. Current studies have shown that Th9 cells play a vital antitumor role in most solid tumors ^25^, but IL-9 has dual roles in tumor immunity. As a lymphocyte growth factor, IL-9 plays a dual role, promoting tumor progression in hematological tumors and inhibiting their development in solid ones, by activating innate or adaptive immune responses. However, these roles are not absolute: in some solid tumors, IL-9 also has the function of promoting tumor development. Finally, the fermentative bacterial products as the short chain fatty acids may impact on colorectal carcinogenesis by favoring the expression of Foxp3 gene, and boosting Tregs’ functions^26 27^.

Given these premises, our study aims to investigate immune system-microbiota crosstalk in CRC through the cellular and molecular characterization of immunity and the comparative evaluation of microbiota composition in healthy and tumor mucosa, focusing on the correlation between the cytokine/chemokine profile and the GM composition in CRC and CRC-associated tissues.

## 2. MATERIAL AND METHODS

### 2.1 Patients recruitment

Forty-five patients affected by non-metastatic colorectal adenocarcinoma at preoperative staging undergoing surgical resection at the Careggi University Hospital (Florence, Italy), were enrolled Specifically, patients’ characteristics are summarized in Table 1. Exclusion criteria were: extraperitoneal rectum localization of the tumor, previous surgery for cancer; previous chemo-radiotherapy treatment; immunodeficiency; trip to exotic areas in the last 5 years; treatment with immunosuppressive drugs, antibiotics or probiotics during the previous 2 months; acute gastrointestinal infections in the month prior to enrollment; concomitant presence of established malignancies or chronic intestinal inflammatory diseases (Crohn’s disease and Ulcerative recto colitis). Tissue pieces of tumor (CRC) and surrounding healthy mucosa (CRC-S) were obtained from the surgical specimen after surgery. The study has received the local Ethics Committee approval (CE: 11166_spe) and an informed written consent has been obtained from each participant.

**Table 1.**
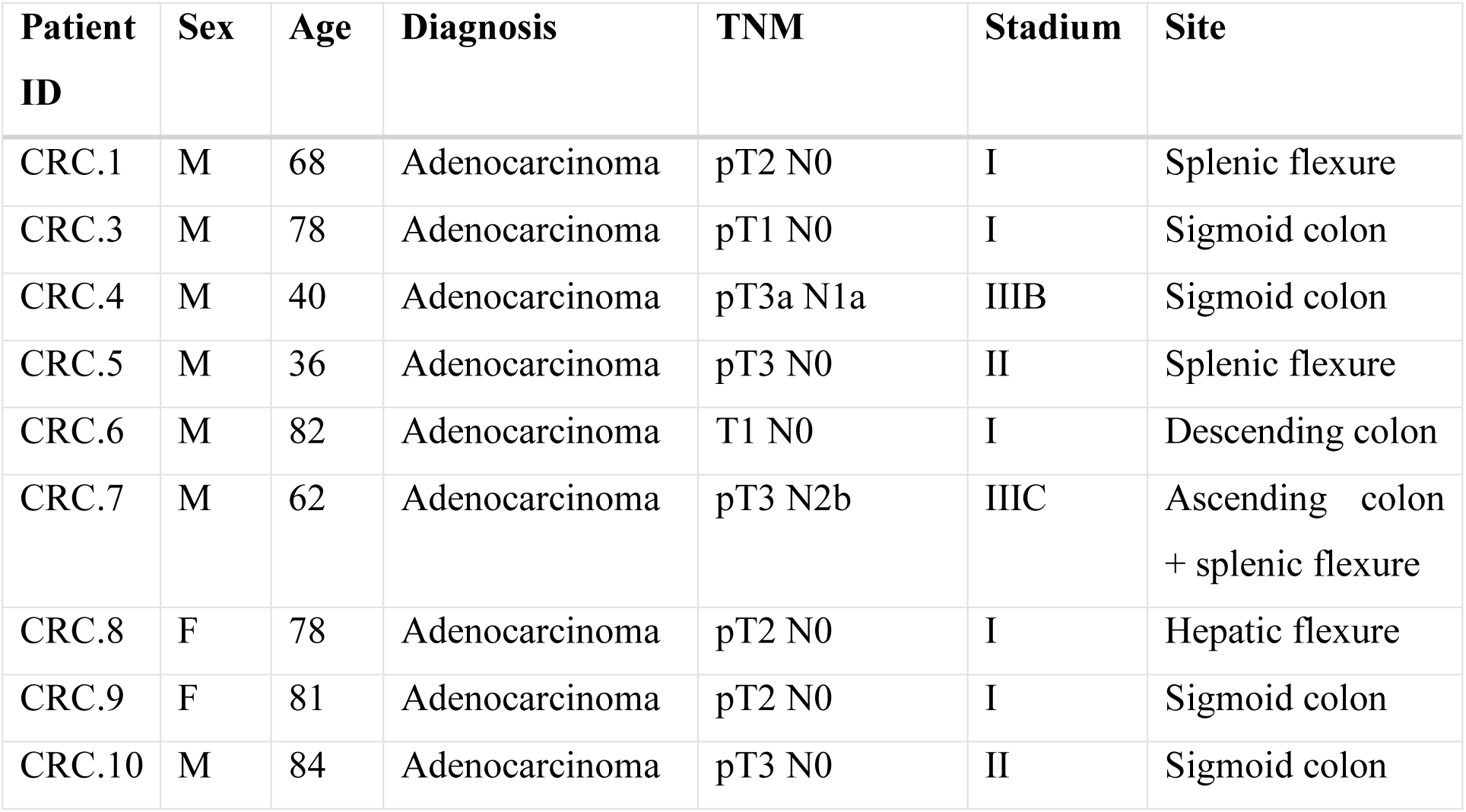
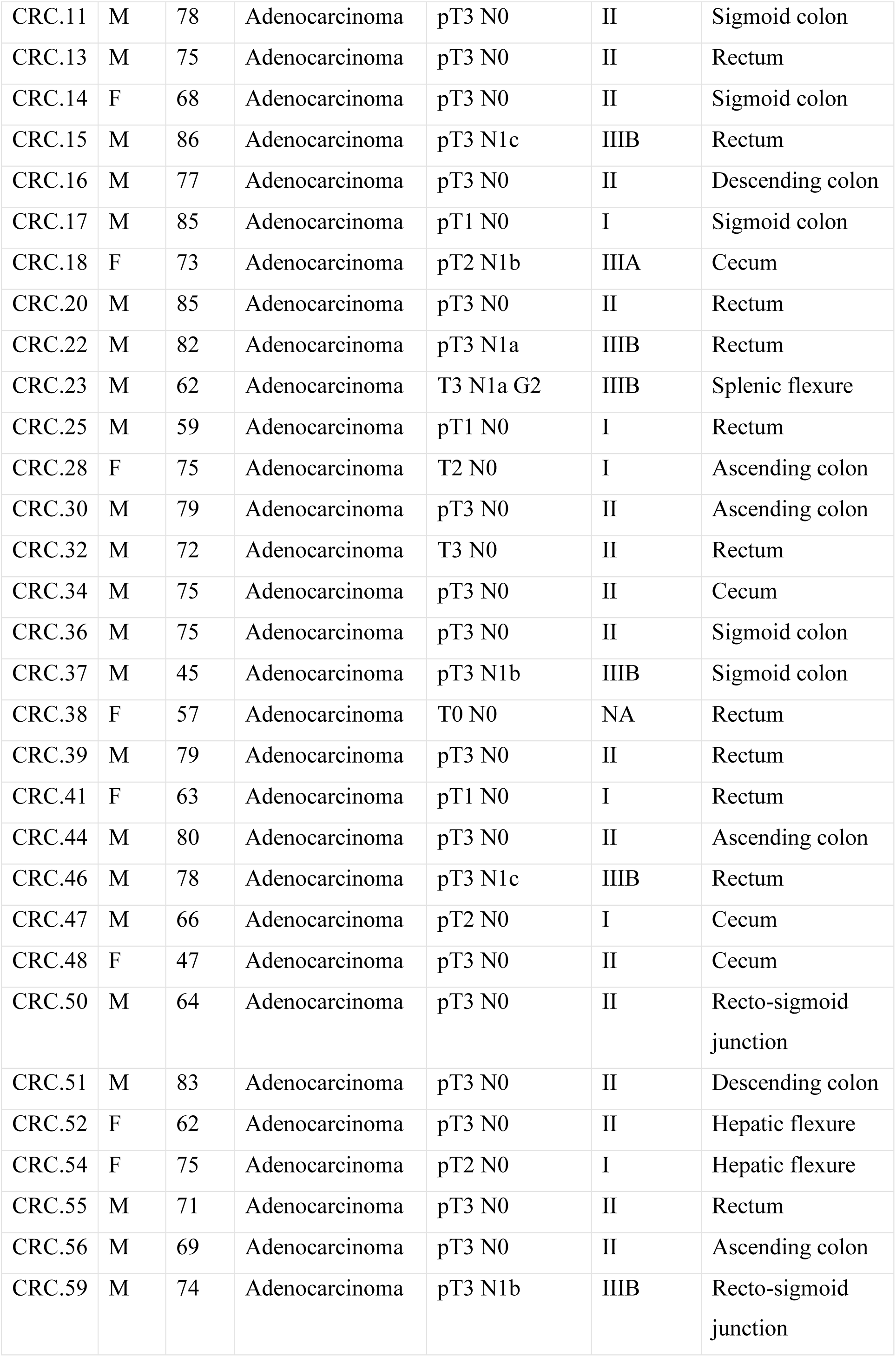
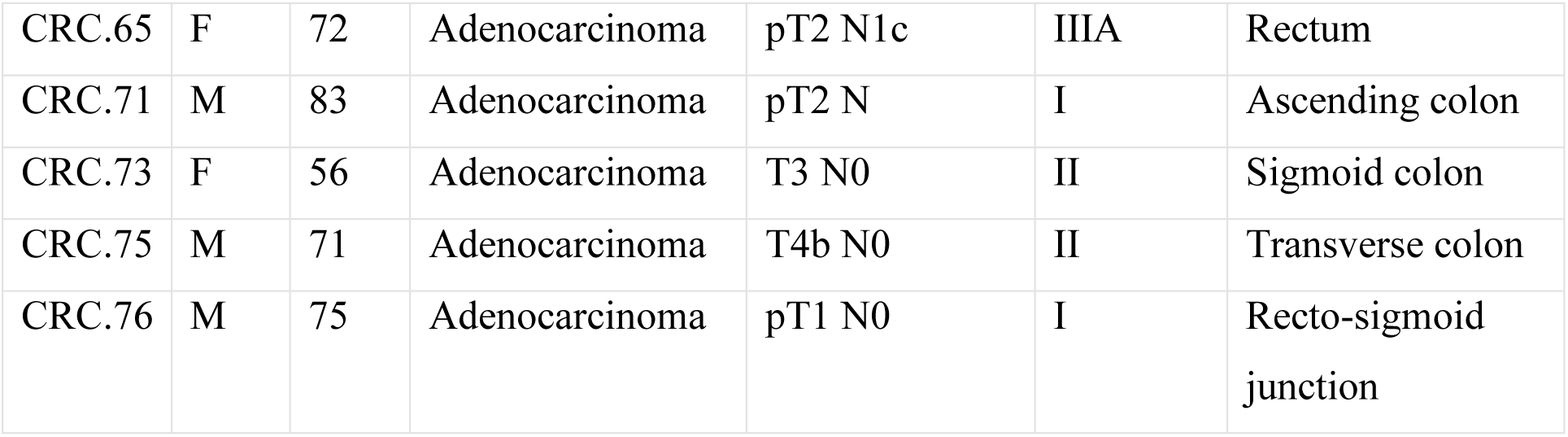
CRC patients’ clinical features.

| Patient ID | Sex | Age | Diagnosis | TNM | Stadium | Site |
| --- | --- | --- | --- | --- | --- | --- |
| CRC.1 | M | 68 | Adenocarcinoma | pT2 N0 | I | Splenic flexure |
| CRC.3 | M | 78 | Adenocarcinoma | pT1 N0 | I | Sigmoid colon |
| CRC.4 | M | 40 | Adenocarcinoma | pT3a N1a | IIIB | Sigmoid colon |
| CRC.5 | M | 36 | Adenocarcinoma | pT3 N0 | II | Splenic flexure |
| CRC.6 | M | 82 | Adenocarcinoma | T1 N0 | I | Descending colon |
| CRC.7 | M | 62 | Adenocarcinoma | pT3 N2b | IIIC | Ascending colon<br>+ splenic flexure |
| CRC.8 | F | 78 | Adenocarcinoma | pT2 N0 | I | Hepatic flexure |
| CRC.9 | F | 81 | Adenocarcinoma | pT2 N0 | I | Sigmoid colon |
| CRC.10 | M | 84 | Adenocarcinoma | pT3 N0 | II | Sigmoid colon |
| CRC.11 | M | 78 | Adenocarcinoma | pT3 N0 | II | Sigmoid colon |
| CRC.13 | M | 75 | Adenocarcinoma | pT3 N0 | II | Rectum |
| CRC.14 | F | 68 | Adenocarcinoma | pT3 N0 | II | Sigmoid colon |
| CRC.15 | M | 86 | Adenocarcinoma | pT3 N1c | IIIB | Rectum |
| CRC.16 | M | 77 | Adenocarcinoma | pT3 N0 | II | Descending colon |
| CRC.17 | M | 85 | Adenocarcinoma | pT1 N0 | I | Sigmoid colon |
| CRC.18 | F | 73 | Adenocarcinoma | pT2 N1b | IIIA | Cecum |
| CRC.20 | M | 85 | Adenocarcinoma | pT3 N0 | II | Rectum |
| CRC.22 | M | 82 | Adenocarcinoma | pT3 N1a | IIIB | Rectum |
| CRC.23 | M | 62 | Adenocarcinoma | T3 N1a G2 | IIIB | Splenic flexure |
| CRC.25 | M | 59 | Adenocarcinoma | pT1 N0 | I | Rectum |
| CRC.28 | F | 75 | Adenocarcinoma | T2 N0 | I | Ascending colon |
| CRC.30 | M | 79 | Adenocarcinoma | pT3 N0 | II | Ascending colon |
| CRC.32 | M | 72 | Adenocarcinoma | T3 N0 | II | Rectum |
| CRC.34 | M | 75 | Adenocarcinoma | pT3 N0 | II | Cecum |
| CRC.36 | M | 75 | Adenocarcinoma | pT3 N0 | II | Sigmoid colon |
| CRC.37 | M | 45 | Adenocarcinoma | pT3 N1b | IIIB | Sigmoid colon |
| CRC.38 | F | 57 | Adenocarcinoma | T0 N0 | NA | Rectum |
| CRC.39 | M | 79 | Adenocarcinoma | pT3 N0 | II | Rectum |
| CRC.41 | F | 63 | Adenocarcinoma | pT1 N0 | I | Rectum |
| CRC.44 | M | 80 | Adenocarcinoma | pT3 N0 | II | Ascending colon |
| CRC.46 | M | 78 | Adenocarcinoma | pT3 N1c | IIIB | Rectum |
| CRC.47 | M | 66 | Adenocarcinoma | pT2 N0 | I | Cecum |
| CRC.48 | F | 47 | Adenocarcinoma | pT3 N0 | II | Cecum |
| CRC.50 | M | 64 | Adenocarcinoma | pT3 N0 | II | Recto-sigmoid<br>junction |
| CRC.51 | M | 83 | Adenocarcinoma | pT3 N0 | II | Descending colon |
| CRC.52 | F | 62 | Adenocarcinoma | pT3 N0 | II | Hepatic flexure |
| CRC.54 | F | 75 | Adenocarcinoma | pT2 N0 | I | Hepatic flexure |
| CRC.55 | M | 71 | Adenocarcinoma | pT3 N0 | II | Rectum |
| CRC.56 | M | 69 | Adenocarcinoma | pT3 N0 | II | Ascending colon |
| CRC.59 | M | 74 | Adenocarcinoma | pT3 N1b | IIIB | Recto-sigmoid<br>junction |
| CRC.65 | F | 72 | Adenocarcinoma | pT2 N1c | IIIA | Rectum |
| CRC.71 | M | 83 | Adenocarcinoma | pT2 N | I | Ascending colon |
| CRC.73 | F | 56 | Adenocarcinoma | T3 N0 | II | Sigmoid colon |
| CRC.75 | M | 71 | Adenocarcinoma | T4b N0 | II | Transverse colon |
| CRC.76 | M | 75 | Adenocarcinoma | pT1 N0 | I | Recto-sigmoid junction |

### 2.2 Immunological analysis

#### 2.2.1 Analysis of tissue infiltrating lymphocytes (TILs)

Tissue pieces were dissociated with the Tumor Dissociation Kit, human (Miltenyi Biotech, UK) in combination with the gentleMACS™ Octo Dissociator (Miltenyi Biotech, GmbH), to obtain a gentle and rapid generation of single-cell suspensions. Then, TILs were magnetically isolated with anti-human CD3 microbeads (Miltenyi Biotech, UK) using AutoMACS Pro Separator (Miltenyi Biotech, GmbH) and analyzed by polychromatic flow cytometry. In detail, TILs from dissociated tissues were characterized for the expression of CD4, CD8, CD25, CD127, IFN-γ, IL-4, IL-17, IL-9, IL-22, and FoxP3 using intracellular cytokine staining. Briefly, TILs were cultured in RPMI 1640 culture medium (SERO-Med GmbH, Wien) supplemented with 10% FCS HyClone (Gibco Laboratories, Grand Island, NY, USA) and stimulated for 5 h using the Leukocyte Activation Cocktail, with BD GolgiPlug™ (BD Pharmingen). Cells were stained for the surface antigens, then fixed with 4% (v/v) paraformaldehyde and permeabilized with 0.5% saponin, followed by intracellular staining with anti-IL-4, anti-IL-17, anti-IL-22, anti-IL-9 and anti-IFN-γ mAbs (BD Biosciences). For the detection of peripheral Tregs, TILs were fixed and permeabilized using the BD Pharmingen Human FoxP3 Buffer Set (BD Biosciences). A minimum of 10,000 events was acquired.

#### 2.2.2 Molecular inflammatory response evaluation

The inflammatory response has been evaluated in homogenized CRC and CRC-S of 14 cancer patients, through specifically assembled kits MixMatch Human 26 Panel for Luminex MAGPIX detection system (Affymetrix, eBioscience) and following the manufacturers’ instructions. In detail, macrophage inflammatory protein-1α (MIP-1α), interleukin (IL)-27, IL-1β, IL-2, IL-4, IL-5, interferon gamma-induced protein 10 (IP-10), IL-6, IL-8, IL-10, IL-12p70, IL-13, IL-17A, granulocyte-macrophage colony stimulating factor (GM-CSF), tumor necrosis factor-α (TNF-α), interferon (IFN)-α, IFN-γ, monocyte chemotactic protein 1(MCP-1), IL-9, P-selectin, IL-1α, IL-23, IL-18, IL-21, soluble intercellular adhesion molecule-1 (sICAP-1) and IL-22 were analyzed. The levels of cytokines were estimated using a 5‐parameter polynomial curve (ProcartaPlex Analyst 1.0). The Low and Upper Limit of Quantification (LLOQ and ULOQ) used for the cytokines and chemokines are reported in Table 2. A value under the LLOQ was considered as 0 pg/ml.

**Table 2.** Low and Upper Limit of Quantification (LLOQ and ULOQ) for each evaluated cytokine/chemokine

| **Cytokine/chemokine** | **ULOQ**<br>**(pg/ml)** | **LLOQ**<br>**(pg/ml)** |
| --- | --- | --- |
| **MIP-1α** | *1880* | *1,84* |
| **IL-27** | *82000* | *20* |
| **IL-1β** | *8250* | *2,01* |
| **IL-2** | *26900* | *6,57* |
| **IL-5** | *30400* | *7,42* |
| **IP-10** | *7750* | *1,89* |
| **IL-6** | *37800* | *9,23* |
| **IL-8** | *9850* | *2,4* |
| **IL-17A** | *9550* | *2,33* |
| **IFN- $\gamma$** | *12675* | *12* |
| **GM-CSF** | *41000* | *10* |
| **TNF-α** | *29500* | *7,2* |
| **MCP-1** | *14800* | *3,61* |
| **IL-9** | *31000* | *7,57* |
| **IL-1α** | *3000* | *0,73* |
| **IL-18** | *49500* | *12* |
| **IL-21** | *39700* | *9,69* |
| **IL-22** | *82500* | *20* |
| **P-selectin** | *5051600* | *1233* |
| **sICAP1** | *870200* | *212* |
| **IL-4** | *36200* | *8,84* |
| **IL-10** | *9250* | *2,26* |
| **IL-12p70** | *28100* | *6,86* |
| **IL-13** | *13400* | *3,27* |
| **IL-23** | *68500* | *17* |
| **IFN-α** | *2250* | *0,55* |

#### 2.2.3 Statistical analysis of immunologic data

Statistical analysis was performed with the SPSS statistical software (version 24). Differences between T cells subsets’ data obtained from CRC and CRC-S samples and tissue cytokines levels evaluated in the same groups were assessed with paired Wilcoxon signed-rank test. P values less than 0.05 were considered statistically significant.

### 2.3 Microbiota characterization

#### 2.3.1 DNA extraction

Genomic DNA was extracted using the DNeasy PowerLyzer PowerSoil Kit (Qiagen, Hilden, Germany) from frozen (−80°C) CRC and CRC-S according to the manufacturer’s instructions. Briefly, tissues were added to a bead beating tube and thorough homogenized with TissueLyser II for 5min at 30 Hz. Total genomic DNA was captured on a silica membrane in a spin column format and subsequently washed and eluted. The quality and quantity of extracted DNA was assessed using the NanoDrop ND-1000 (Thermo Fisher Scientific, WalthAP, US) and the Qubit Fluorometer (Thermo Fisher Scientific), respectively. Then, genomic DNA was frozen at −20°C.

#### 2.3.2 Bioinformatic analysis of 16S rRNA

Extracted DNA samples were sent to IGA Technology Services (Udine, Italy) where amplicons of the variable V3–V4 region of the bacterial 16s rRNA gene were sequenced in paired-end (2 ×300 cycles) on the Illumina MiSeq platform, according to the Illumina 16S Metagenomic Sequencing Library Preparation protocol^28^.

Raw sequences were processed following the software pipeline MICCA^29^. Paired end reads were assembled using the “mergepairs” command, maintaining a minimum overlap of 100 bp and an edit distance in the maximum overlap of 32 bp. Subsequently the sequences were cut with the “trim” command in order to remove the primers and eventually eliminate the reads with imperfect primers sequences. All the reads with a length lower than 350 bp and with an error rate higher than or equal to 0.5 were removed with the “filter” command.

Cleaned reads were eventually merged into a single file with the “merge” command and transformed into a fasta file. The OTUs were generated using the “otu” command in “denovo_greedy” mode, setting a 97% identity and performing an automatic removal of chimeras with the “-c” option. The longest sequence of each OTU was used for the taxonomic assignment using the “classify” command in “rdp” mode, i.e. using the RDP Bayesian classifier that is able to obtain classification and confidence for taxonomic ranks up to genus.

#### 2.3.3 Statistical analysis of bacterial communities

Statistical analyses on the bacterial community were performed in R (R Core Team, 2014) with the help of the packages phyloseq 1.26.1^30^, DESeq2 1.22.2^31^, breakaway 4.6.16^32 33^ and other packages satisfying their dependencies, in particular vegan 2.5-5^34^ Rarefaction analysis on OTUs was performed using the function rarecurve (step 50 reads), further processed to highlight saturated samples (arbitrarily defined as saturated samples with a final slope in the rarefaction curve with an increment in OTU number per reads < 1e-5).

For the cluster analysis (complete clustering on euclidean distance) of the entire community the OTU table was first normalized using the total OTU counts of each sample and then adjusted using square root transformation.

The coverage was calculated by Good’s estimator^35^ using the formula: (1 - n/N) × 100, where n is the number of sequences found once in a sample (singletons), and N is the total number of sequences in that sample.

Richness, Shannon, Chao 1 and Evenness indices were used to estimate bacterial diversity in each sample using the function estimate_richness from phyloseq^30^. The evenness index^36^ was calculated using the formula E = S/log(R), where S is the Shannon diversity index and R is the number of OTUs in the sample. Differences in all indices between CRC and CRC-S were tested using a paired Wilcoxon signed-rank test. Sample richness was further measured using the estimator and its associated error introduced in the breakaway package^32^. The function betta_random of breakway was further used to evaluate the statistical differences in richness between paired-by-patient CRC and CRC-S samples.

The differential analysis of abundance at the OTUs as well as at the different taxonomic ranks (created using the tax_glom function in phyloseq) was performed with DESeq2^31^ using a two group blocked by patient design in order to perform a paired test.

### 2.5 Statistical analysis of the association between tissue microbiota and cytokines

The association between tissue microbiota and cytokines was investigated with a 2-step analysis, separately for the mucosa and tumor tissues. In the first step, we implemented a modified version of the Sure Independence Screening (SIS) Procedure^37^. SIS uses the notion of marginal correlation, in our case the correlation of a single cytokine with the dependent variable, to rank the cytokines. The cytokines with smallest p-value from a Dirichlet regression^38^, with that given cytokine as the only predictor, are included in step 2. For each cytokine, the p-value is obtained testing the model with the considered cytokine and the intercept against a model with only intercept with a likelihood-ratio test. Step 1 is necessary only when the list of cytokines is too long with respect to the sample size; in our analysis, we selected the three most relevant cytokines from step 1.

In the second step, we used a Dirichlet Multinomial Regression to determine the joint effect of cytokines on the tissue microbiota. We implemented a Bayesian Variable Selection (BVS) method based on a thresholding function^39^. This approach is based on a Monte Carlo Markov Chain algorithm that explores the space of possible models. The method’s output is a list of posterior probability of inclusion (PPI) and the posterior mean of the non-zero regression coefficients. PPI is the probability, between 0 and 1, that a given association cytokine-genera is non-zero, accounting for the effect all other cytokines. The posterior mean is an estimate of a non-zero association. Each estimated regression coefficient evaluates the taxon-cytokine association, whose sign and magnitude measure the effect of the cytokine on the taxon.

## 3. RESULTS

### 3.1 Assessment of tissue infiltrating T cell subsets’ distribution in healthy and cancer mucosa

We performed the polychromatic flow cytometry analysis of TILs’ isolated from the dissociated CRC and CRC-S. The percentage of CD4^+^ and CD8^+^ TILs in the mucosa samples group did not differ significantly. In detail, the mean percentages (SD) of CD4^+^ cells were 58.03 (6.88) in CRC vs 58.15 (6.32) in CRC-S and the mean percentages of CD8^+^ T cells were 17.42 (5.65) in CRC vs 14.39 (4.14) in CRC-S.

The analysis of the T cells subsets revealed that the tumor mucosa samples group displayed higher percentages of Th17 (CRC vs CRC-S: 10.02 (4,32) vs 5.13 (1.39); p=0,0008), Th2 (CRC vs CRC-S: 3.45 (1.31) vs 1.41 (0.93); p=0,0011) and Treg (CRC vs CRC-S: 4.08 (1.44) vs 2.10 (0.57); p=0,0040) as shown in Figure 1A. Regarding the T cytotoxic cells, the CRC group showed higher percentages of Tc17 (CRC vs CRC-S: 6.33 (3.98) vs 1.77 (1.00), p=0,0036), Tc1/Tc17 (CRC vs CRC-S: 6.25 (4.74) vs 1.88 (1.48), p=0,0022), and Tcreg (CRC vs CRC-S: 1.08 (0.81) vs 0.06 (0.08), p=0,0055) (Figure 1B). Notably the number of Th9 is major (but not significant) in CRC tissue while the Tc9 are similar in the two different sites.

**Figure 1.**
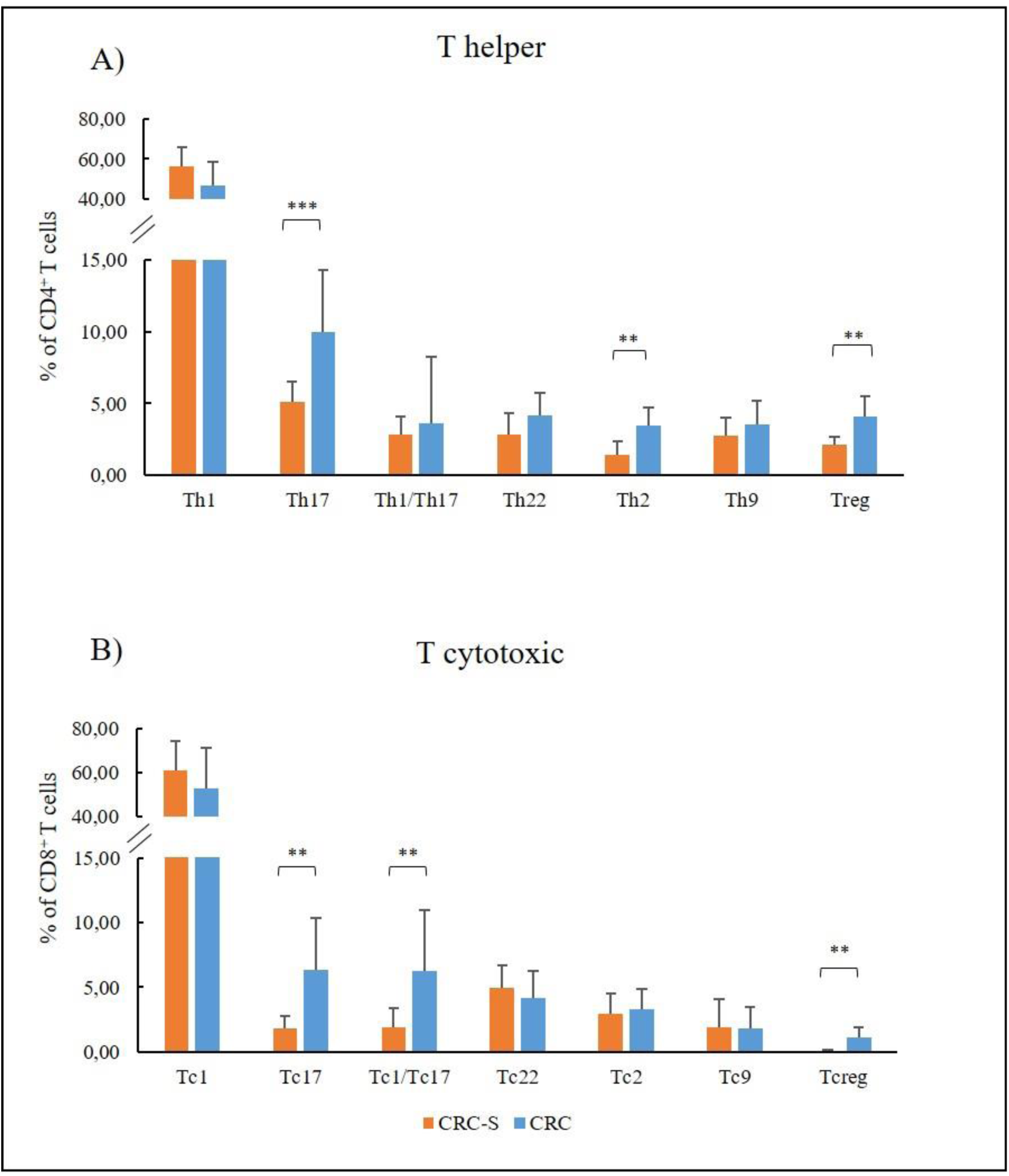
T cells subsets distribution in the tumor mucosa (CRC) and surrounding healthy mucosa (CRC-S) samples groups. Panel **A**) reports the mean percentages (^+^SD) of T helper subsets in respect to the percentage of CD4^+^ T cells, while panel **B**) reports the mean percentages (^+^SD) of T cytotoxic subsets in respect to the percentage of CD8^+^ T cells. Statistical analyses were calculated using Wilcoxon signed-rank test. The asterisks (*) represent p-values, *p < 0.05, **p < 0.01, ***p < 0.001.

### 3.2 Molecular inflammatory profile in CRC-associated tissues

We compared the molecular inflammatory profile of the homogenized CRC and CRC-S of 14 cancer patients, through the evaluation of 26 pro- and anti-inflammatory cytokines. Six of the evaluated cytokines (IL-4, IL-10, IL12p70, IL-13, IL-23 e IFN-α) were under the LLOQ (Table 2) in all samples, either because levels were very low (not detectable) or because these molecules are not produced. The other 20 cytokines showed a common trend in all patients, characterized by higher levels in CRC compared to CRC-S (Figure 2A). In particular, MIP-1α, IL-1β, IL-2, IP-10, IL-6, IL-8, IL-17A, IFN-γ, TNF-α, MCP-1, IL-9, IL-1α, P-selectin, were increased significantly in CRC compared to CRC-S (Figure 2B).

**Figure 2.**
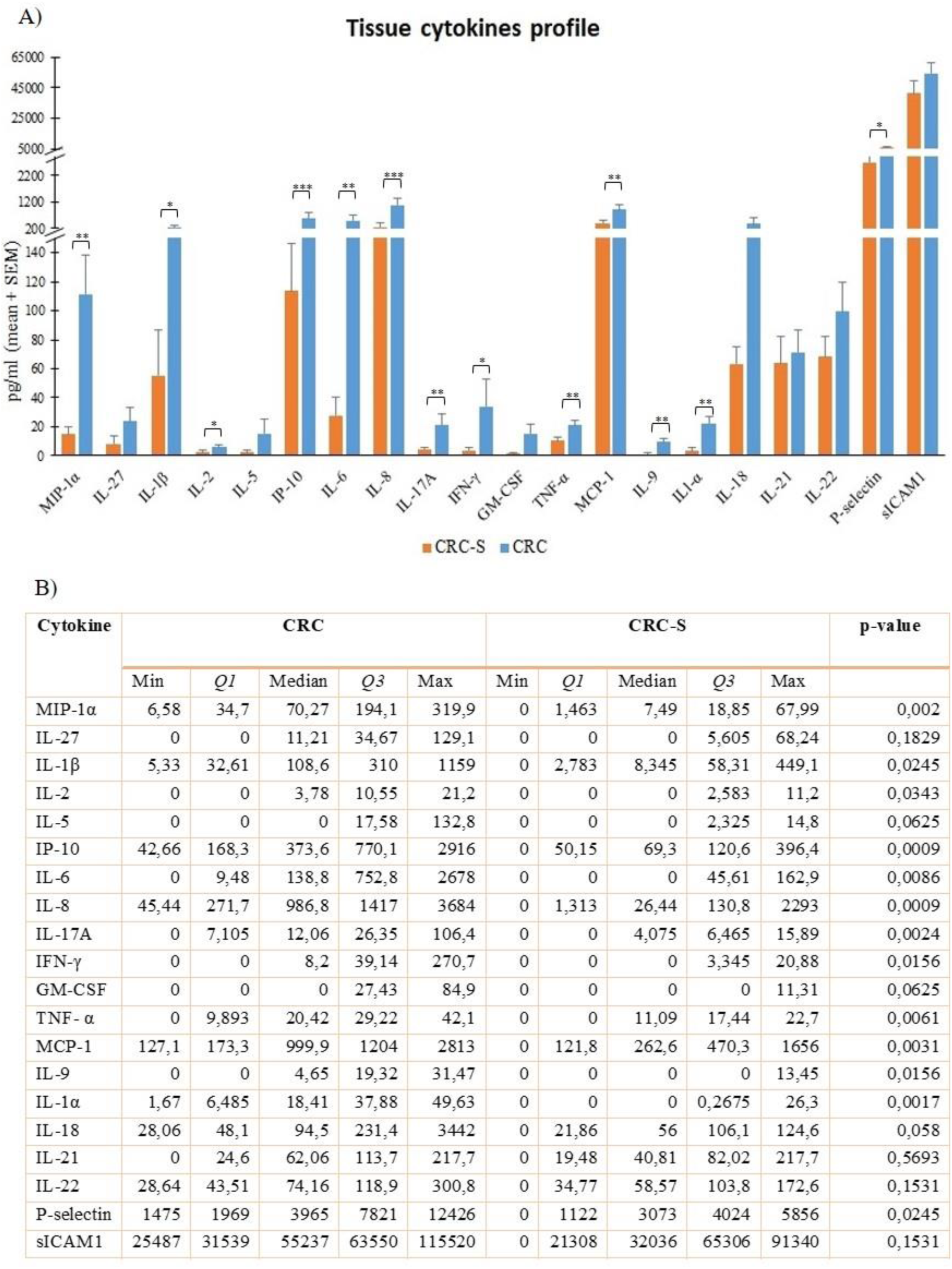
Tissue cytokine levels in 14 CRC patients. **A**) Histogram reports the mean (+ SEM) cytokines levels (pg/ml) of the evaluated cytokines in CRC-S and CRC of 14 CRC patients. **B**) Statistical report of the evaluated cytokines in CRC-S and CRC tissues. Wilcoxon signed rank test was performed to test the differences between CRC-S and CRC paired samples. A p-value < 0,05 is considered statistically significant. The asterisks (*) represent p-values, *p < 0.05, **p < 0.01, ***p < 0.001. CRC-S= Healthy mucosa; CRC= tumor mucosa; Q1= 1st quartile; Q3= 3rd quartile.

### 3.3 Comparison of mucosal microbiota composition in CRC and CRC-S

Our sequencing efforts in assessing microbiota composition encompassed a total of 12,475,251 reads for 40 sample pairs. After all the steps of pre-processing, that included pair merging, trimming, quality filtering and chimera detection, a total of 8,458,126 (67.8 %) were available for further analysis. As shown in Figure 3, saturation curves revealed that most specimens were sufficiently sampled. Samples showed a Good’s coverage ranging from 99 to 100% indicating that less than 1% of the reads in a given sample came from OTUs that appear only once in that sample.

**Figure 3.**
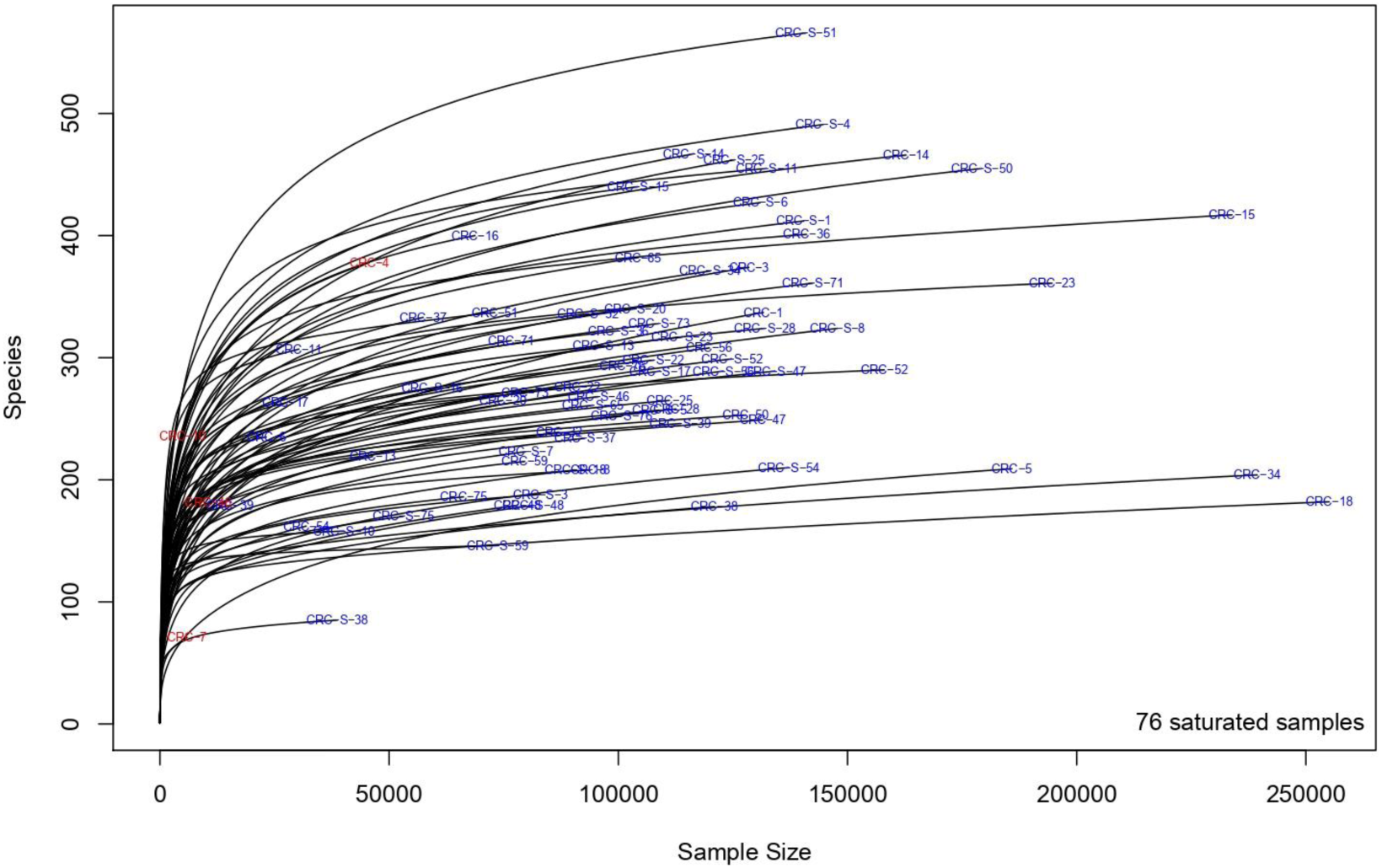
Rarefaction curves showing the level of saturation of OTUs.

As shown in Figure 4, the alpha diversity of samples did not display significant differences for Shannon index and Evenness. On the contrary, a significant (p = 0.011) Chao1 index evidenced that rare OTUs are enriched in CRC-S vs CRC, denoting a higher diversity.

**Figure 4.**
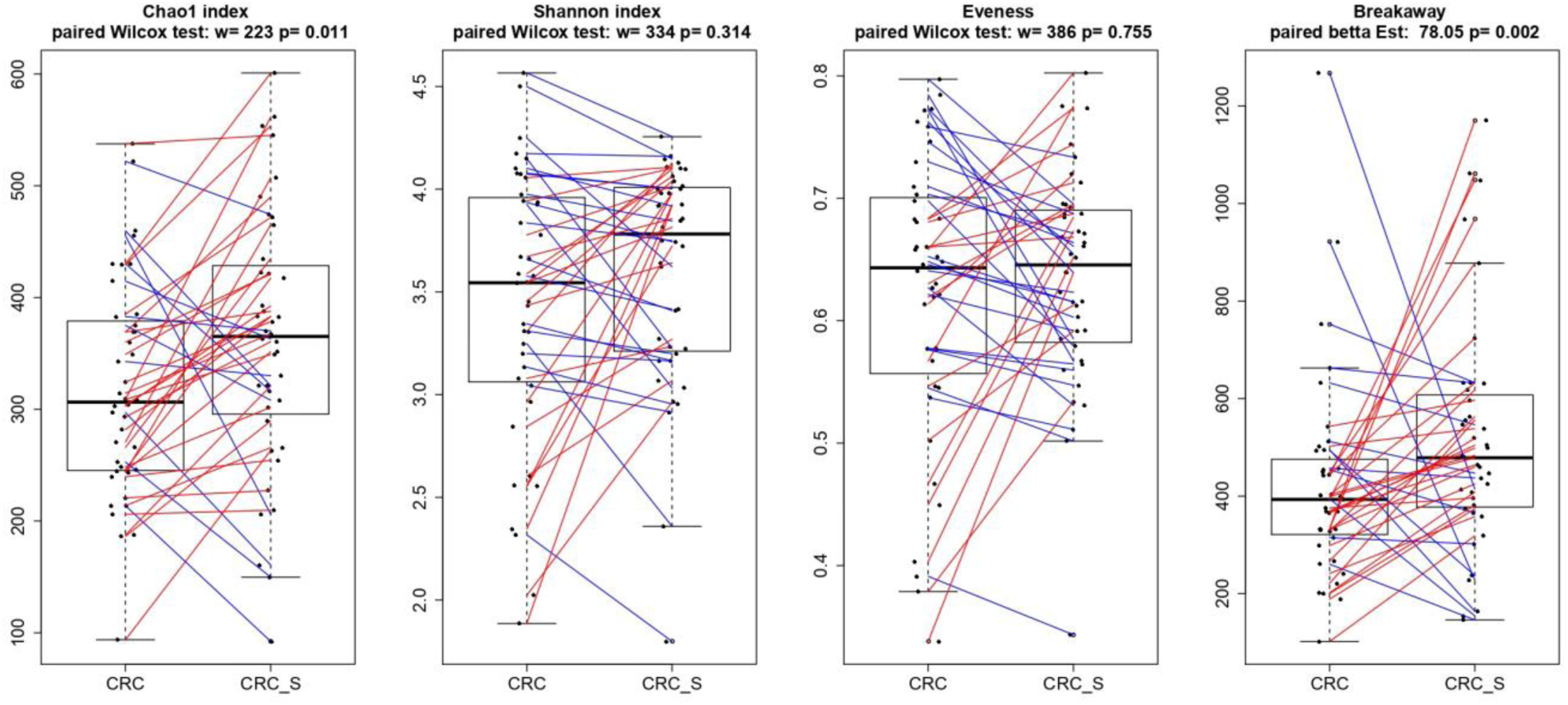
Box-plots illustrating alpha diversity indices (Chao1 index, Shannon index, Evenness and Breakaway) in CRC and CRC-S samples. Statistical differences were evaluated using paired Wilcoxon signed-rank test for Chao, Shannon and Evenness indices and using the paired betta analysis implemented in the Breakaway R package. P-values less than 0.05 were considered statistically significant.

Taxonomic analysis detailed in Table 3 revealed for the 2454 OTUs formed the confident (<20% error) presence of 29 phyla (>99% reads), 50 classes (>98% reads), 87 orders (>98% of reads), 176 families (>96% reads) and 372 genera (>86% reads). Stacked boxplots of taxa abundance at different taxonomic ranks are available in **Supplementary Figure S1**.

**Table 3.**
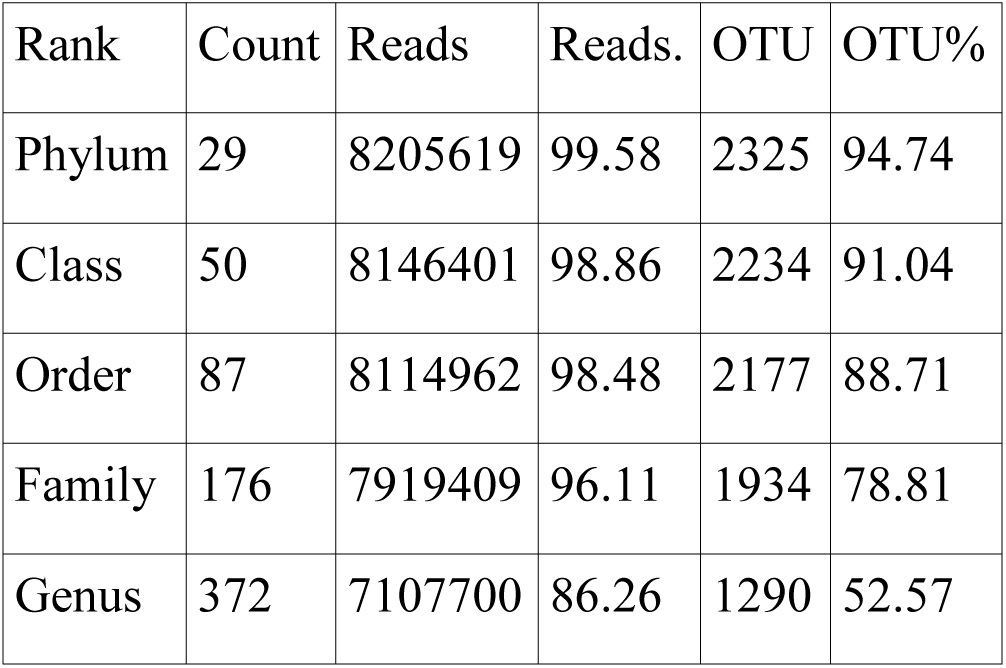
Summary of the taxonomic analysis of the obtained OTUs.

In order to investigate and confirm the paired nature of sampling (i.e. tumor tissue vs. surrounding healthy tissue) we performed a cluster analysis on normalized OTU counts. As shown in Figure 5, we verified that 36/40 paired samples were in effect paired also in terms of microbial composition, a result robust to changes in distance metrics (e.g. Bray-Curtis) and clustering method (data not shown).

**Figure 5.**
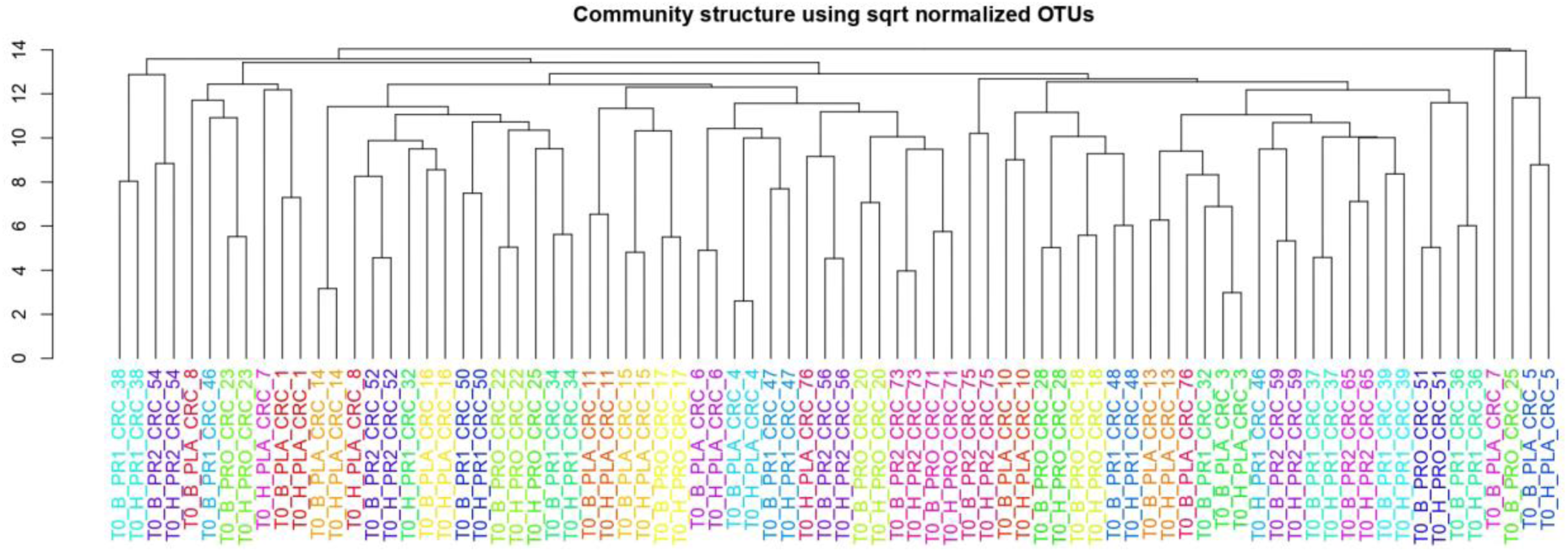
Cluster analysis on normalized OTU counts.

The analysis of the taxonomic composition reported that 6 phyla dominated the dataset (98% sequences), namely *Firmicutes* (41.24%), *Bacteroidetes* (35.89%), *Proteobacteria* (13.37%), *Fusobacteria* (4.68%), *Verrucomicrobia* (2.10%) and *Actinobacteria* (1.56%) as shown in Figure 6.

**Figure 6.**
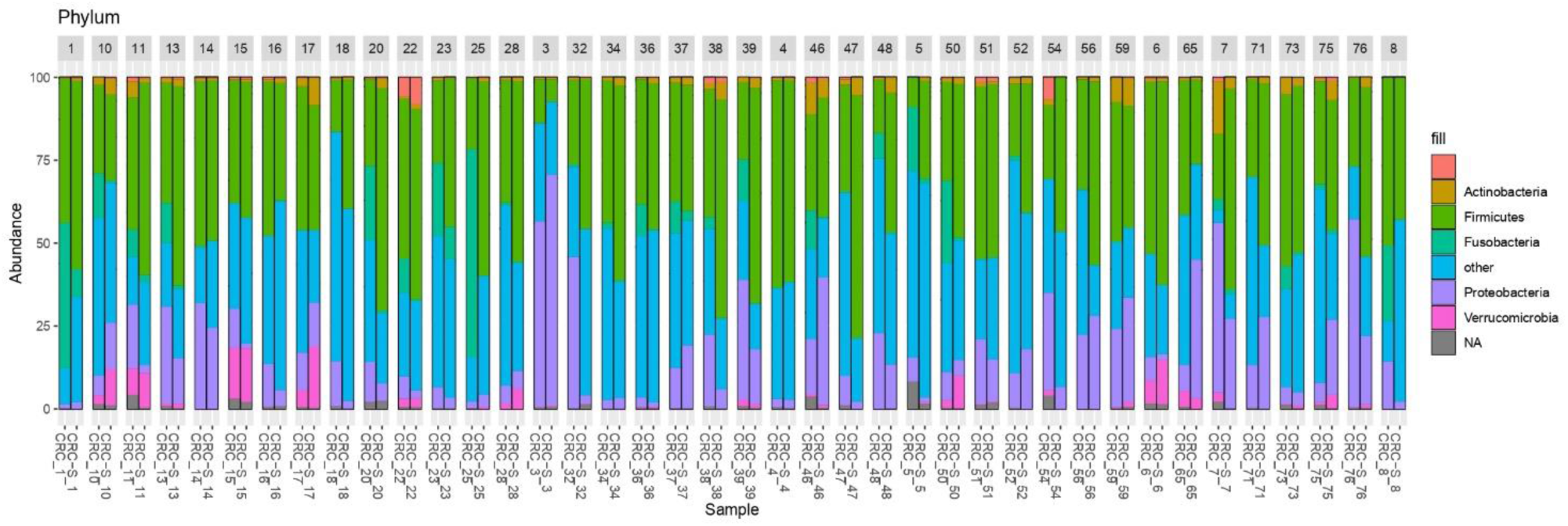
Stacked boxplots of microbial composition at phylum level of CRC and CRC-S samples.

The paired comparison of the abundance of single OTU revealed significant (adj.p<0.05, abs (logFC)>=1) differences between CRC and CRC-S samples groups, with 6.6% OTUs involved. At the phylum level *Fusobacteria* and *Proteobacteria* were significantly higher in CRC compared to CRC-S (logFC = −2.92, adj.p = 6.17e-15 and logFC = −0.95, adj.p = 1.92e-05, respectively). At the genus level 14.7% genera were observed as significantly (adj.p<0.05, abs (logFC)>=1) different, the most abundant being *Fusobacterium* (average OTUs 6199, log2FC = −2.93, adj.p 1.06e-08), *Ruminococcus2* (*Lachnospiraceae* family, average OTUs 2911, log2FC = 1.31, adj.p = 1.38e-3) and *Ruminococcus* (*Ruminococcaceae* family, average OTUs 1640, log2FC = 1.82, adj.p = 4.12e-05). The complete results of the differential analysis at all taxonomic ranks is available in supplementary data **S2**.

### 3.4 Correlation of the cytokine profile with the microbiota composition in CRC-associated tissues

To evaluate this critical and crucial point, we first applied the screening procedure SIS (step 1). We considered the OTU counts of the three more abundant genera and aggregated all other genera in a residual category. Our dependent variable was then defined by four categories: *Bacteroides spp.*, *Prevotella spp.* and *Escherichia/Shigella spp.* (plus the residual category). The four top-ranked cytokines were IL-18, IFN-γ, IL-5 and IL-2. Using these four cytokines (plus the intercept), we ran the Bayesian Variable Selection method (step 2). In the tumor tissues, we detected the association between IL-5 and *Prevotella spp.*, with a PPI=0.81; the same association was found to be supported by the data collected from the mucosa tissues as well (PPI=0.77). The effect of IL-5 on *Prevotella* is estimated to be positive and equal to 0.64 (posterior mean) and 0.86, for tumor and mucosa tissues, respectively. However, this analysis hardly describes the entire picture since many cytokines with very small p-values in the first step were not included in the second step due to computational constrains.

Moreover, we decided to perform a second analysis that includes directly into step 2 the cytokines that showed differential abundances in comparative analysis between CRC and CRC-S samples. However, due to the computational constrains that impose a limit on the sample size; we selected some of the cytokines that were significantly increased in CRC compared to CRC-S samples, according to their relevance in the current literature. In particular, we have chosen IFN-γ, IL-17A, IL-8, IL-1β, IL-1α, IP-10, MIP-1α and IL-9. The results of the BVS Dirichlet Multinomial Regression (step 2) are reported in Table 4 and Table 5.

**Table 4.** Posterior probabilities of inclusion (PPIs).

|  | *Prevotella*<br>*spp.* | *Bacteroides*<br>*spp.* | *Escherichia/Shigella*<br>*spp.* | Residual<br>category |
| --- | --- | --- | --- | --- |
| intercept | 1.00 | 1.00 | 1.00 | 1.00 |
| IFN-γ | 0.10 | 0.00 | 0.45 | 0.00 |
| IL-17A | 0.82 | 0.00 | 0.00 | 0.00 |
| IL-8 | 0.00 | 0.00 | 0.00 | 0.00 |
| IL-1β | 0.00 | 0.00 | 0.00 | 0.00 |
| IL-1α | 0.00 | 0.00 | 0.00 | 0.00 |
| MIP-1α | 0.00 | 0.00 | 0.00 | 0.00 |
| IP-10 | 0.00 | 0.00 | 0.77 | 0.00 |
| IL-9 | 0.95 | 0.83 | 0.00 | 0.00 |

**Table 5.** Posterior mean of the regression coefficients.

|  | *Prevotella*<br>*spp.* | *Bacteroides*<br>*spp.* | *Escherichia/Shigella*<br>*spp.* | Residual<br>category |
| --- | --- | --- | --- | --- |
| intercept | -1.10 | 0.41 | -0.93 | 1.90 |
| IFN-γ | -0.05 | 0.00 | 0.31 | 0.00 |
| IL-17A | -1.08 | 0.00 | 0.00 | 0.00 |
| IL-8 | 0.00 | 0.00 | 0.00 | 0.00 |
| IL-1β | 0.00 | 0.00 | 0.00 | 0.00 |
| IL-1α | 0.00 | 0.00 | 0.00 | 0.00 |
| MIP-1α | 0.00 | 0.00 | 0.00 | 0.00 |
| IP-10 | 0.00 | 0.00 | -0.89 | 0.00 |
| IL-9 | 1.37 | -0.91 | 0.00 | 0.00 |

From the results, we can notice that *Prevotella spp.* is associated with both IL-17A and IL-9. The first association is negative, the effect is 1.08 (posterior mean), whereas the second is positive, posterior mean of 1.37. *Bacteroides spp.* and *Escherichia/Shigella spp.* show a negative association with IL-9 and IP-10 respectively, posterior mean equal to −0.91 and −0.89 respectively.

## 4 DISCUSSION

In this study, we explored the immune responses, the microbiota and their association in colorectal cancer, comparing the composition of T cell subsets, the cytokine/chemokine pattern and the gut microbiota composition in tumoral and surrounding mucosa sample of CRC patients. First, the TIL’s subsets assessment revealed higher percentages of Th17, Th2, Th9 and Tregs, together with increased percentages of Tc17, Tc1/Tc17 and Tcregs, in CRC samples compared to CRC-S. These data confirm with our previous findings, where we documented the weakening of the effector functions of T cell clones isolated from adjacent healthy mucosa compared to the cancerous one and the increase of T lymphocytes’ subsets (Th2/Th0/Tregs/Tnull) that can promote tumor progression^9^. Contextually, the Th2 harmful role in tumor development has been well established; a Th2 shift in the tumor microenvironment, especially for CRC, strongly contributes to cancer relapse, metastasis, and worse prognosis^40 41 42^. Moreover, although current evidence for the role of Th17, Th9 and Tregs in cancer are contradictory, excess inflammation caused by CD4^+^ and CD8^+^ IL-17 producing T cells or the immunosuppression induced by Treg may lead to carcinogenesis^43 44^.

Tumor infiltrating Th17 and Tc17 cells have been found in various human cancers, confirming their pro-tumorigenic properties^45 46 47^. In particular recent finding suggests pivotal role of Th17 in the CRC pathogenesis, as they trigger and amplify local inflammatory processes and tumorigenesis^42 48 49^. According to our results, different studies found higher percentages of Th17 cells in tumor tissues compared to adjacent non-tumor tissues, in both CRC and other malignancies^50 51 52 53^. Also, Zhang et al. demonstrated that, compared with normal tissues, Tc17 cells specifically accumulated in tumor tissues of cervical tumor patients^54^, while Mielke et al. found a significant enrichment of Tc17 genes in CRC samples, indicating that these T cell subset are involved in CRC development^55^. In addition, according to Kuang et al., which reported high levels of Tc17 cells in hepatocellular carcinoma patients^56^, we found a significant increase of Tc1/Tc17 cells in CRC samples. Hence, the current literature indicated that tumor environment promote enrollment of Tc1/Tc17 cells that endowed with a pro-inflammatory effect, but an attenuated cytotoxic potential.

Concerning CD4^+^ and CD8^+^ Treg cells, their role in prevention of immune hypersensitivity and extensive inflammatory responses is strongly established. However, through their immunosuppressive properties, Tregs can favor immune escape mechanisms of tumor cells and that is why high amounts of peripheral or tumor-infiltrating CD4^+^ or CD8^+^ Tregs are often associated with poor clinical outcome in various types of cancer^57 58 59 60^. According to our findings, high number of tumor infiltrating CD4^+^ and CD8^+^ Tregs, compared to surrounding non-tumor tissue, is widely documented^58 61 62 63 64 65^.

Finally, among these subpopulations, Th9 cells are relatively new, and less is known about their role on tumor immunity, especially in CRC development.

Interestingly, IL-4 and TGF-β are requested for the differentiation of naïve T cells into Th9^66^. Moreover, TGF-β can promote the transformation of Th2 cells into Th9 cells^67^, and IL-4 can transform Tregs into a Th9 cell subset^68^. Moreover, together with IL-4, IL-1β sustain the differentiation of Th9 cells^69^. Usually, the Th9 cells, activating both the innate and adaptive immune responses, play an important role in antitumor immunity, such as documented in melanoma and lung cancer^70 71^. In addition, recent studies have shown that also Tc9 cells have a stronger antitumor effect than conventional IFN-γ-producing CD8^+^ T cells, in the same cancers (melanoma and breast cancer)^72 73^. Anyway, as a lymphocyte growth factor, IL-9 can promote the lymphocytes activation and proliferation, exerting tumorigenic role in different hematological tumors^74^. It’s tumor promoting ability has been demonstrated also in solid cancers. In line with our findings, Tan and collaborators observed that the number of Th9 cells was significantly increased in human hepatocellular carcinoma (HCC) tissue. Moreover, they demonstrated their tumour promoting role in HCC via the CCL20 and STAT3 signalling pathways and their correlation with poor patient survival rate^75^.

Besides the TILs analysis, although many studies investigated the levels of several plasma cytokines in CRC patients^76 77 78 79^, we assessed for the first time the molecular inflammatory profile of homogenized CRC and CRC-S samples through the evaluation of an exhaustive custom panel of 26 (pro- and anti-inflammatory) cytokines/chemokines. Although six cytokines were undetectable or not produced, the other 20 were all increased in CRC compared to the CRC-S. In particular, CRC samples revealed significant higher levels of MIP-1α, IL-1β, IL-2, IP-10, IL-6, IL-8, IL-17A, IFN-γ, TNF-α, MCP-1, IL-1α, P-selectin and IL-9.

First, the relevant higher percentages of chemokines that we detected in CRC samples, such as MCP-1, MIP-1α, IL-8 and IP-10, reflects a colonic inflammation. Indeed, many studies demonstrated their role in the development of a tumor-favoring microenvironment due to their abilities to favor angiogenesis and to stimulate macrophages and CD8^+^ T cells recruitment *in situ^80 81 82 83 84^*. According to our results, different studies found high levels of these chemokines in colon cancer tissues compared to healthy tissues^85 86 87^.

As regards pro-inflammatory cytokines like IL-1α, IL-1β, IL-6, IL-17Aand TNF-α, the evidences support their pro-tumorigenic role because of their ability to promote tumor initiation, progression, angiogenesis and metastasis in many human malignancies, including the CRC^88 89 90 91^. Then, our findings are consistent with previous studies^92 93^ that indicated an increased tissue and serum expression of these cytokines in CRC patients compared to healthy controls^94 95 96^.

On the contrary, although the overexpression of IFN-γ in CRC tissues compared to normal ones^97^ can be considered positive because of its established robust antitumor activity ^98 99^, IL-2 and IL-9 displayed both pro-tumor and anti-tumor potentials^100 101^.

IL-2, despite its pleiotropic effects on immune system, has been approved for the treatment of metastatic renal cell carcinoma and metastatic melanoma^102^. Moreover, an immunotherapy strategy with IL-2 successfully demonstrated a reduction of the tumor growth in a mouse colon cancer model *in vivo*^*103*^. However, in line with our results, high levels of IL-2 in CRC patients was observed also by Bosek et al.^104^. Regarding IL-9, contrary to our result, Wang et al. shown that IL −9 was less expressed in human colon carcinoma^105^. However, even Huang et al. reported low amounts of IL-9 in tissue and plasma of CRC patients but these low levels were associated with tumor progression^106^. Interestingly, Tian and colleagues have shown that the expression of IL-9 in colitis-associated cancer (CAC) tissue was significantly higher than that in adjacent tissues and the Lentiviral vector-mediated IL-9 overexpression in the colon cancer cells lines RKO and Caco2 could promote their proliferation^107^. Moreover, Carlsson documented higher level of IL-9 in the serum of breast cancer patients and its correlation with tumor metastasis^108^.

Furthermore, since the relationship between the origin and development of CRC and the imbalance of GM has been well established during past years^109 110 111^, we characterized the composition of tumor and adjacent healthy mucosa. First, we verified that 36/40 paired samples were in effect paired also in terms of microbial composition and these data confirms and further highlight the intrinsic individual nature of GM^112^. Then we found that alpha diversity of CRC and CRC-S samples did not displayed significant differences for Shannon index and Evenness. However, according to Yang et al. and Liu et al., a significant Chao1 index evidenced that rare OTUs are enriched in CRC-S vs CRC^113 114^.

Since it is well established that *Fusobacteria* and *Fusobacterium spp.* are associated with CRC and are amplified during colorectal carcinogenesis^115,116 117^, the obtained data are in agreement with the current literature and with our previous results^118^. In addition, we found a significantly higher percentage of *Proteobacteria* in CRC compared to CRC-S, according to evidence that an imbalanced GM often is associated with a sustained increase in *Proteobacteria* phylum members^119 120^. Consistent with Weir et al.^121^ we found that *Ruminococcus spp.* were more represented in CRC compared to CRC-S samples, though some authors reported that in patients with colon carcinoma *Ruminococcus spp.* have low prevalence compared to normal population^122 123^.

Finally, in order to explore the interplay among the intestinal microbiota and local immune response in CRC patients, for the first time we correlated the cytokines’ profile and the intestinal microbiota composition of CRC and CRC-S samples using the BVS Dirichlet Multinomial Regression. Firstly, we applied the screening procedure SIS and considering the OTU counts of the three more abundant genera (*Bacteroides spp., Prevotella spp.* and *Escherichia/Shigella spp* and aggregated all other genera in a residual category), the four top-ranked cytokines were IL-18, IFN-γ, IL-5 and IL-2, which we used to run the Bayesian Variable Selection method. Although these analyses cannot adequately describe the complex scenario of the relationship between secreted cytokines and intestinal composition, we observed the positive association between IL-5 and *Prevotella spp.*, both in tumor and mucosa tissues.

As well known, IL-5 is essential for eosinophil differentiation/survival; it is produced by Th2 lymphocytes and mast cells and is associated with several allergic diseases including allergic rhinitis and asthma^124 125^. Eosinophilia has also been observed in cancer, including breast, ovarian, cervical, prostate cancers and CRC but the prognosis link remains controversial and different for tumor types. Eosinophil infiltration is considered unfavorable in Hodgkin’s lymphoma, but positive in breast and prostate cancers^126^. As recently reviewed^127^, higher numbers of infiltrating eosinophils detected in CRC tissue were repeatedly shown to be prognostically favorable^128 129 130^, but the mechanisms of CRC growth inhibition remain poorly understood.

In vitro studies assessing tumoricidal activity of eosinophils against CRC lines, have shown that the effect was mediated by eosinophil-produced TNF-α and granzyme A and that the eosinophil attachment to cancer cells depended on interaction LFA-1 and ICAM-1 that was upregulated by IL-18^131 132^. In addition, a recent CRC mouse model found that tumor-homing eosinophils secrete chemoattractants for CD8^+^ effector T cells, eventually causing tumor rejection^133^.

Finally, Hollande et al found that IL-33 expressed by murine tumor cells induced eotaxin 1 production that led to eosinophil recruitment and degranulation-dependent suppression of tumor growth^134^. In addition, IL-33 is a potent eosinophil activator stimulating their degranulation and it was reported that IL-33-deficient mice had GM dysbiosis and were highly susceptible to both colitis and colitis-associated cancer^135^, which could also be related to impaired eosinophil-driven responses.

Moreover, abundant IL-5 levels are documented in the synovium of rheumatoid arthritis patients^136^ and notably Scher et al. showed that the presence of *Prevotella copri* was strongly correlated to rheumatoid arthritis^137^. So, taking into account all these reported data, in addition to our results that showed a positive correlation between IL-5 and *Prevotella spp.* and given the anti-inflammatory role of IL-5, we could assume an attempt to restore an eubiotic ecosystem in the colon mucosa contrasting the CRC development.

However, when it comes to the immune-microbiota crosstalk, we always have to keep in mind that selected members of the same genus have different disease modulating properties in different diseases. Recently, Calcinotto et al.^138^ demonstrated in a transgenic mouse that *Prevotella heparinolytica* promotes the differentiation of Th17 cells colonizing the gut and migrating to the bone marrow, where they favour progression of multiple myeloma. In addition, they observed that the treatment with antibodies blocking IL-17 and, of note, IL-5, reduced the accumulation of Th17 cells and eosinophils and delayed disease progression, suggesting a positive correlation of *Prevotella spp.* and IL-5, in agreement with our results.

Subsequently, we performed a second analysis including in the Bayesian Variable Selection method all cytokines that showed differential abundances in the comparative analysis of CRC and CRC-S: IFN-γ, IL-17A, IL-8, IL-1β, IL-1α, IP-10, MIP-1α and IL-9. We only focused on tumor tissues, because the same analysis performed on data from the mucosa tissues did not select any associations. For the first time, we found that always *Prevotella spp.* is negatively associated with IL-17A but positively related to IL-9. In addition, *Bacteroides spp.* and *Escherichia/Shigella spp.* shown a negative association with IL-9 and IP-10 respectively.

Although we found a negative correlation between *Prevotella spp.* and IL-17A, it has recently been discovered that *Prevotellaceae* are able to promote Th17 cells differentiation and *Prevotella spp*. are associated with Th17-mediated diseases, including periodontitis and rheumatoid arthritis^138 139^. These conflicting data could be explained by speculating that the same bacterial taxa in different environmental conditions, as previously declared, may have different roles, even contrasting ones. However, we cannot exclude that contrasting results may derive from the relatively low level of taxonomic resolution of 16S rRNA gene metagenomics, which cannot fully discriminate between different species of the same genus (as *Prevotella spp*.). We can hypothesize that different *Prevotella* species colonize different human body districts. Indeed, the pangenome of *Prevotella* spp. is large, indicating high genomic diversity of the different species^140^ as well as a high differentiation among strains of the same species, as for instance *P. copri^141^*. It is well documented that bacteria can modulate the lymphocyte differentiation by the SCFAs (as previously^142^), whose composition can vary (qualitatively and quantitatively) based on the inflammatory status, as we have recently shown for different intestinal gastrointestinal pathologies, such as celiac disease, adenomatous polyposis and CRC^143^. Maybe the periodontal *Prevotella spp.* related to Th17 cells stimulation are not subdued to a dysbiotic environment such as the inflamed colon cancer mucosa. In addition, in a study in which *Prevotella histicola* was used to modulate immune response and treat arthritis in a humanized mouse model, Marietta et al. reported that treated mice showed significantly lower levels of IL-17, in agreement with our data. The authors reported also decreased level of IL-9 as compared to placebo treated mice, in contrasting with our results ^144^. Finally, Campisciano^145^ detected an increase of the relative vaginal abundance of *Prevotella timonensis* in women infected with HPV, which showed a decreased concentration of the IL-15, IL-7, and IL-9, that however they associated with the virus infections.

Regarding the negative correlation between *Bacteroides spp.* and IL-9, we have not found studies documenting this association but since it is well know that the Th17 produces the IL-9, we can have indirect evidence of this correlation. Round et al. found that *Bacteroides fragilis* inhibits Th17 development, inducing Tregs’ accumulation and Vaahtovuo et al. demonstrated the lower abundance of *Bacteroides spp.* in rheumatoid arthritis patients^146 147^.

Our results also showed that *Escherichia/Shigella spp.* are negative correlated with IP-10 that is induced in many viral, bacterial and parasite infections, i.e. shigellosis an *E. coli* infection^148 149^. Because IP-10 resulted increased in CRC samples, also the negative correlation between *Escherichia/Shigella spp.* can be supported.

To conclude, our data describe a clear dissimilarity of the cellular and molecular inflammatory profile and intestinal microbiota composition between the tumor and the adjacent healthy tissue, displaying the generation of a peculiar CRC microenvironment. In particular, the increased percentage of both CD4^+^ and CD8^+^ T cells proving a strong reaction of the immune system against the cancer. However, the presence *in situ* of Tregs and T CD8^+^ effector cells testify the effort of the immune response but also its double-edged role because just these lymphocytes are involved in the enhancing of cancer progression. Furthermore, the higher percentages of several cytokines/chemokines produced in the tumor tissue homogenate demonstrated that different types of immune cells are involved in these complex anticancer responses although some of these may even encourage the neoplastic progression. In addition, the correlation of the cytokine profile with the intestinal microbiota composition in CRC and CRC-S samples confirmed the presence of a bidirectional crosstalk between the immune response and the host’s commensal microorganisms.

Finally, our data showed for the first time that *Prevotella* and *Bacteroides* species are correlated positively and negatively, respectively, with the IL-9 in human CRC. We are aware that other studies in humans and in animal models will be needed to strengthen our data, but there is no doubt regarding the innovativeness of our study. We, for the first time have highlighted a relationship between the intestinal microbiota and the presence of IL-9 that usually identify the Th9, one of the new T lymphocytes’ subsets, whose role in tumor development, also for CRC, is still debated.

