## Supplemental figures for "Significant and conflicting correlation of IL-9 with *Prevotella* and *Bacteroides* in human colorectal cancer"

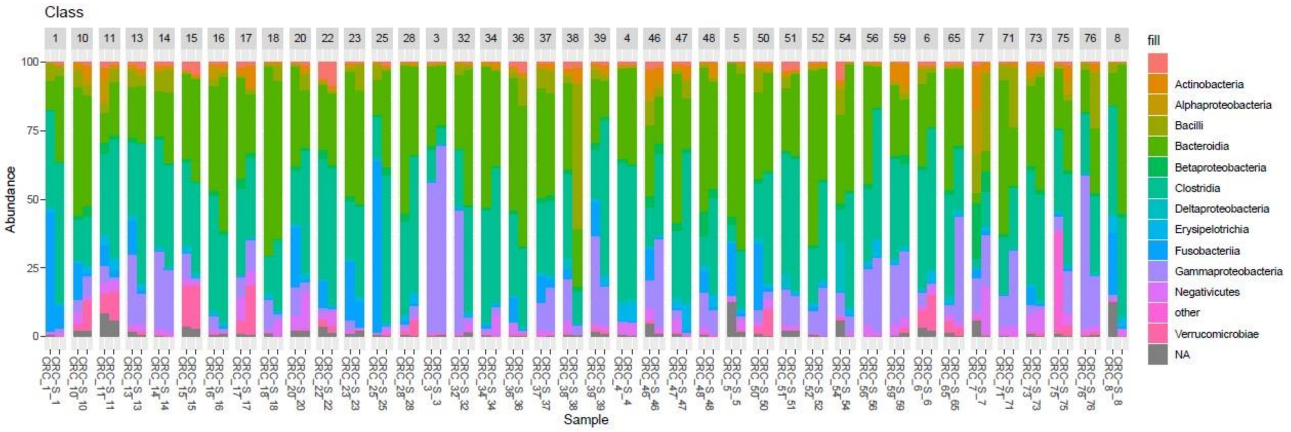


**Figure S1.** Stacked boxplots of microbial composition at class level of CRC and CRC-S samples.

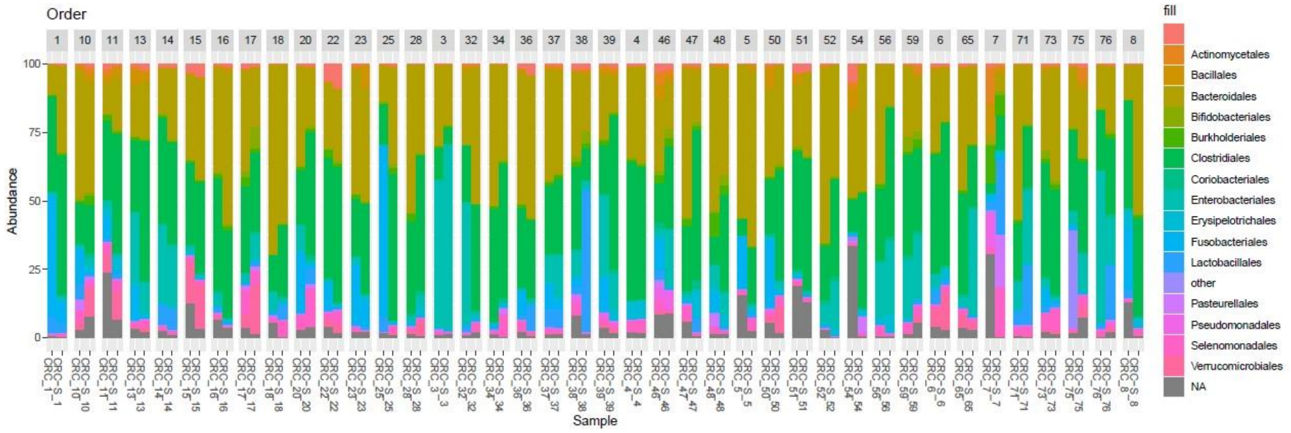


**Figure S2.** Stacked boxplots of microbial composition at order level of CRC and CRC-S samples.

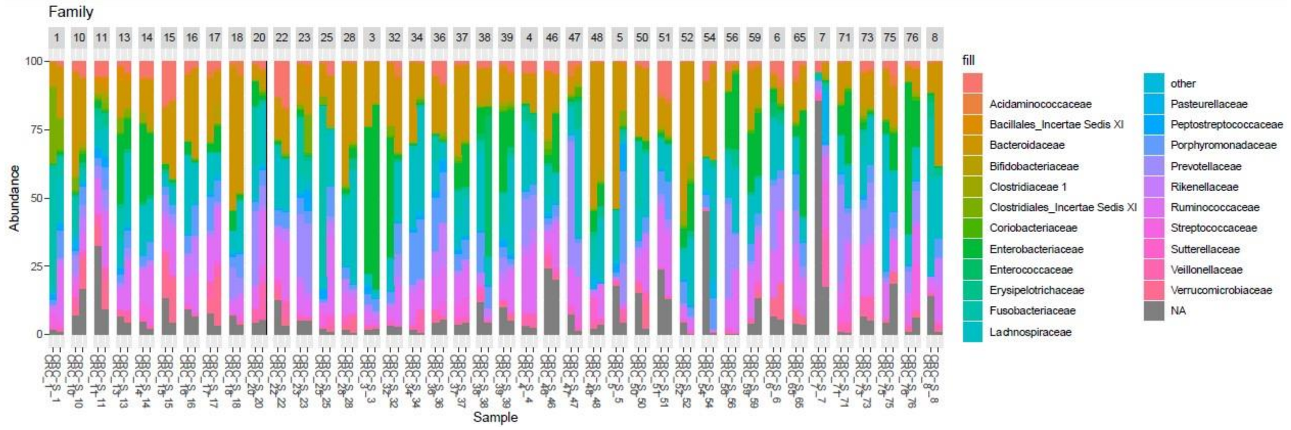


**Figure S3.** Stacked boxplots of microbial composition at family level of CRC and CRC-S samples.

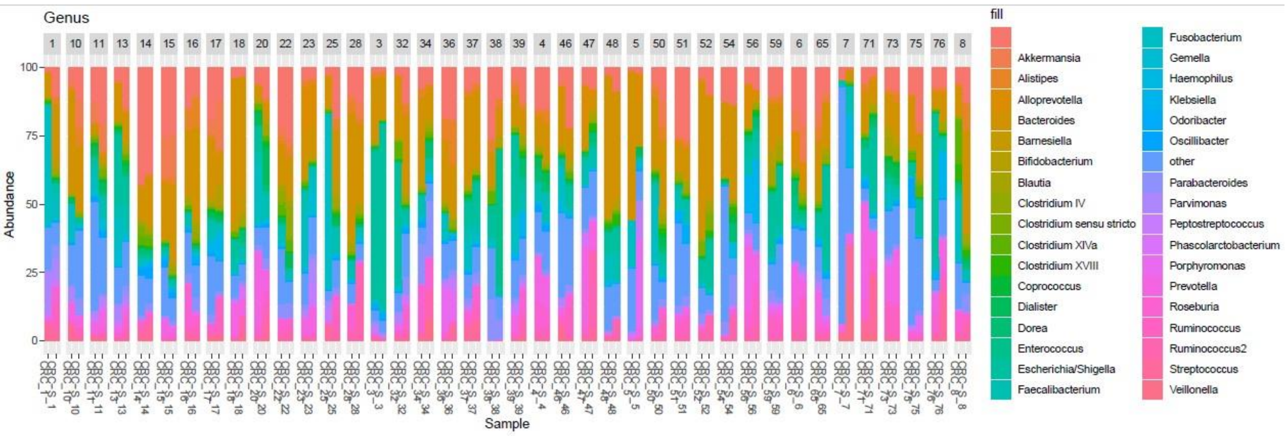


**Figure S4.** Stacked boxplots of microbial composition at genus level of CRC and CRC-S samples.
